# Mixed-species interactions constrain diversification and shape biofilm evolution

**DOI:** 10.64898/2026.02.26.708127

**Authors:** Sanaa Harrass, Joyce To, Parisa Noorian, Gustavo Espinoza-Vergara, Vaughn S. Cooper, Scott A. Rice, Diane McDougald, M. Mozammel Hoque

**Affiliations:** Australian Institute for Microbiology & Infection, University of Technology Sydney, Sydney, NSW, Australia; Center for Evolutionary Biology and Medicine, University of Pittsburgh, School of Medicine, Pittsburgh, PA, USA; School of Civil, Mining and Environmental Engineering, University of Wollongong, NSW, Australia

**Keywords:** Biofilm, LTEE, diversity, evolution, phenotypes, mutation

## Abstract

Long-term experimental evolution (LTEE) provides a powerful framework for dissecting how ecological interactions shape adaptive trajectories. Here, we evolved *Klebsiella pneumoniae, Pseudomonas protegens* and *Pseudomonas aeruginosa* in single- and mixed-species biofilm communities for 24 weeks and tracked changes in population dynamics, phenotypes, and genomes. In mono-species evolution, all three species exhibited similar dynamics of adaptation, with steadily increasing biofilm-associated populations. In contrast, mixed-species communities displayed striking compositional shifts, with *P. protegens* emerging as the dominant biofilm former and *K. pneumoniae* dominating the supernatant. Phenotypic assays revealed that all three species showed enhanced biofilm formation, but this increase was consistently greater in isolates from mono-species than mixed species communities, with *P. protegens* showing the largest gains. Beyond biofilm production, biofilm-associated isolates exhibited greater phenotypic diversification than planktonic isolates, whereas mixed-species interactions constrained diversification. Whole-genome sequencing identified species-specific putative adaptations such as *csrD* in *K. pneumoniae, yfiBNR* in *P. protegens*, and *cheA* in *P. aeruginosa* that arose early, persisted, and were enriched in mixed-species isolates. Functional assays confirmed that these mutations were indeed adaptive by enhancing biofilm formation, with *yfiBNR* mutations in *P. protegens* increasing cyclic-di-GMP production and producing a competitive advantage that recapitulated its dominance in LTEE biofilms. Our findings show that biofilm evolution fosters phenotypic diversification, whereas interspecific interactions shape adaptive trajectories, with specific mutations acting as keystone drivers of long-term ecological dynamics in multi-species communities.

## Introduction

Microbial communities rarely exist as single-species populations in nature; instead, they persist in complex assemblages where interactions among taxa profoundly shape evolutionary trajectories, ecological stability, and functional outputs [1, 2]. Biofilms are surface-associated microbial aggregates embedded in extracellular polymeric matrices and represent a dominant lifestyle for many bacteria in natural, industrial, and clinical environments [3, 4]. This lifestyle confers increased tolerance to environmental stresses and antimicrobials, but also imposes spatial and resource constraints that intensify both competitive and cooperative interactions [5, 6]. Such interactions can influence not only community composition but also the direction and rate of adaptive evolution [7, 8].

Long-term experimental evolution (LTEE) has been instrumental in revealing fundamental principles of microbial adaptation [9, 10]. While classic LTEE studies have focused predominantly on single-species populations grown under well-mixed conditions, there is growing recognition that community context, particularly in structured habitats like biofilms can alter selective pressures and constrain or diversify evolutionary outcomes [11]. Recent work demonstrates that interspecific interactions can promote coexistence through niche partitioning, yet can also limit phenotypic diversification by restricting available adaptive pathways [12, 13]. How these forces play out over hundreds of generations in spatially structured multispecies systems remains poorly understood.

The Gram-negative bacteria *Klebsiella pneumoniae, Pseudomonas protegens* and *Pseudomonas aeruginosa* are ecologically and clinically relevant species that frequently coexist in biofilms across diverse environments, including plant rhizospheres, water systems, and human-associated microbiomes [14, 15]. These species differ in metabolic capabilities, motility and exoproduct repertoires, which can mediate competitive and cooperative behaviors [6, 16]. In mixed-species biofilms, such traits influence spatial positioning, resource capture and community stability [8, 17]. However, whether these differences lead to predictable patterns of dominance, diversification, and genetic adaptation over evolutionary timescales remains unclear.

Here, we used a 24-week LTEE to compare evolutionary and ecological dynamics in mono-species and mixed-species biofilm communities of *K. pneumoniae, P. protegens* and *P. aeruginosa*. By integrating temporal population quantification, high-throughput phenotyping and whole-genome sequencing of 574 isolates, we identified contrasting trajectories of phenotypic diversification, biofilm adaptation, and mutational targets in mono versus mixed-species contexts. Our findings reveal that while biofilm-associated evolution generally increases biofilm formation capacity, interspecific interactions constrain phenotypic heterogeneity, channel adaptation toward specific genetic pathways, and drive long-term shifts in species dominance. These results highlight how a small number of keystone mutations can couple phenotypic adaptation to community-level ecological outcomes in structured multispecies environments.

## Materials and methods

### Bacterial strains and design of LTEE

The wild-type strains of eGFP-tagged *K. pneumoniae* KP-1, eCFP-tagged *P. protegens* Pf-5 (ATCC BAA-477) and mCherry-tagged *P. aeruginosa* PAO1 (ATCC BAA-47) were used to establish single-species and mixed-species communities [6, 18]. LTEE was performed using bead transfer model under biofilm-promoting conditions using sterile polystyrene beads as attachment surfaces according to Poltak and Cooper 2011 [2]. Three and six replicate lineages were maintained for mono and mixed-species communities, respectively. M9 minimal medium supplemented with casamino acid and glucose referred to here as MCG (48 mM Na_2_HPO_4_; 22 mM KH_2_PO_4_; 9 mM NaCl; 19 mM NH_4_Cl; 2 mM MgSO_4_; 0.1 mM CaCl2; 0.04% wt/vol glucose, and 0.2% w/v casamino acids) were used to conduct the LTEE. The assay was conducted in round-bottom, borosilicate glass tube using a rotary shaker at 60 rpm at 24^0^ C. Beads were transferred every three days into 2 mL of fresh medium containing new sterile beads to propagate biofilm-associated populations. Communities were evolved for 24 weeks, with sampling performed weekly.

### Quantification of bead-associated and planktonic populations

The bead-associated and planktonic populations were quantified every week. During, serial passaging of the beads one bead was removed from each lineage into a tube containing sterile 0.9% NaCl for quantification. The tube containing the bead was sonicated in an ultrasonic water bath (Powersonic 420, Thermoline Scientific) at medium power for 1 min, followed by vortexing vigorously to dislodge biofilm-associated cells. Planktonic populations were quantified by removing 100 µL of the supernatant from each lineage. Serial dilutions were plated on CHROMagar media to quantify colony-forming units (CFU) for each species. For the mixed-species communities, CHROMagar allowed differentiation of *K. pneumoniae* (greenish colonies) from *Pseudomonas* spp. (white colonies) based on colony morphology. *P. aeruginosa* were quantified by incubating one CHROMagar plates at 37^0^ C as *P. protegens* do not grow at this temperature. While *P. protegens* were quantified by using Streptomycin (50 µg/mL) into the CHROMagar medium incubated at 30^0^ C.

### Phenotypic assays

A total of 648 single isolates (bead-associated and planktonic, across 4 time points and lineages) were profiled for biofilm formation (Supplementary Table 1). *Pseudomonas* spp. were subjected to motility, protease activity, and pyoverdine production assay. In addition, *P. aeruginosa* was also tested for pyocyanin production. Biofilm formation was quantified using a 96-well crystal violet assay. Overnight cultures were normalized to OD_600_ = 0.05 in MCG, incubated statically at 24°C for 24 h, stained with 0.1% crystal violet, washed and solubilized in 95% ethanol. Absorbance was measured at 590 nm. Motility assays were performed on 0.3% agar (swimming) LB plates, inoculated centrally with 1 μL of overnight culture and incubated at 30°C for 24 h. Motility was quantified as the diameter of colony expansion. Protease activity was assessed on LB agar supplemented with 2% skim milk. Zone diameters of casein degradation were measured after 24 h at 30°C. Siderophore production was measured as pyoverdine fluorescence (excitation 400 nm, emission 460 nm) from culture supernatants normalized to OD_600_, and pyocyanin production in *P. aeruginosa* was quantified spectrophotometrically at 520 nm following separation of the supernatants.

### Sequencing and genomic analysis

Genomic DNA was extracted from all 648 phenotyped isolates using the QIAamp 96 DNA QIAcube HT Kit (Qiagen, cat. no. 51331) according to manufacturer’s instructions. DNA concentration was measured using the Quant-iT™ PicoGreen™ dsDNA Reagent (Invitrogen). Of the 648 isolates, 588 passed the sequencing quality requirements and were subjected to downstream analyses. Whole-genome sequencing libraries were prepared according to Hackflex method [19] and sequenced on Illumina NovaSeq6000 platform (2×150 bp paired-end). Whole-population sequencing was conducted on pooled bead-associated and supernatant fractions from each lineage (three mono and six mixed-species) across four time points. Reads were quality-trimmed using TrimGalore to remove adapter contamination and low-quality bases (≤Q30) and were checked using FastQC. The sequence reads for each of the three species from whole populations samples of mixed-species communities were separated using “bowtie2” followed by “samtools”. Filtered reads were mapped to the respective curated ancestral reference genomes and genetic variants were identified using breseq pipeline [20]. The curated (mutated) ancestral reference genome for the three species were built using “gdtools” in breseq pipeline using the *K. pneumoniae* KP-1 (NZ_CP012883), *P. protegens* Pf-5 (NC_004129) and *P. aeruginosa* PAO1 (NC_002516). Genetic variants in single isolates were identified using clonal mode while whole population samples were identified with using polymorphism mode with polymorphism frequency cut off of 0.1. A sufficient number of reads were obtained from whole populations samples of mixed-species communities. However, data analysis could not be performed for several mixed-species whole-population samples due to insufficient sequencing reads. Specifically, this was the case for *P. aeruginosa* at weeks 8, 15, and 24, and for the *P. protegens* week 1 bead sample.

### Protein structural modelling

Protein structure of the CsrD (A0A3R7K3W9), YfiB (A0A2C9EG30) and CheA (G3XCT6) were obtained from AlphaFold protein structure database (https://www.alphafold.ebi.ac.uk) [21]. Structures were visualized and rendered with the MolStar on AlphaFold website. The amino acid residues were manually denoted and colored in pink according to the respective nsSNPs or deletions observed on CsrD, YfiB and CheA. Alphafold predication model color codes different regions of the protein based on the prediction confidence score (Predicted Local Distance Difference Test: pLDDT). Dark blue (pLDDT > 90) regions indicate very high, cyan (90 > pLDDT > 70) indicate high, yellow/orange (70 > pLDDT > 50) and red (pLDDT < 70) indicate low confidence prediction.

### Cyclic-di-GMP quantification

Quantification of c-di-GMP levels was performed using the c-di-GMP ELISA kit by Cayman Chemical, USA (Item #501780). Briefly, bacterial strains were cultured in MCG for overnight, normalized to OD600=0.4 and 1mL of the normalized culture pelleted for lysis in 80uL of B-PER™ buffer (Thermo Fisher Scientific). Cell lysates were serially diluted for the assay according to manufacturer’s instructions. Concentration of c-di-GMP in the samples were calculated from the standard curve, readings outside of the recommended range and inconsistent concentrations (after dilution factor adjustment) were excluded. Corresponding sample dilution factors were multiplied to obtain the final c-di-GMP concentrations in picograms per milliliter.

### Competition assays

Competition experiments were conducted in LTEE-mimicking conditions using wild-type ancestral strains or isolates carrying prevalent mutations in *csrD* (*K. pneumoniae*), *yfiBNR* (*P. protegens*) or *cheA* (*P. aeruginosa*). Equal initial densities of each competitor were inoculated into 2 mL of MCG medium containing sterile polystyrene beads. Tubes were incubated at 24^0^C at 60 rpm on rotary shaker. Bead-associated and planktonic fractions were sampled after 24 h. Relative abundances were determined by plating on CHROMagar and calculating the proportion of CFUs for each species as described earlier.

### Statistical analysis

Statistical analyses were performed in R v4.3. Data were log_10_-transformed where necessary to meet normality assumptions. Biofilm formation, motility, and other phenotypic measures were compared between mono and mixed-species isolates using Kruskal-Wallis tests. Principal component analysis (PCA) was performed using the “princomp” function on phenotypic data and visualized using “factoextra” package in R. Mutation frequencies over time were visualized with ggplot2. Significance thresholds were set at *P* < 0.05.

## Results

### LTEE reveals contrasting population dynamics in mono and mixed-species biofilm communities

We conducted a 24-week LTEE experiment in which *K. pneumoniae, P. protegens* and *P. aeruginosa* were propagated either as single species or as a three-species community under repeated bead-transfer biofilm selection (Fig. 1). At each transfer, colonized beads were moved into fresh medium containing a new sterile bead, imposing a strong and recurrent population bottleneck while maintaining continuous opportunities for biofilm growth and dispersal. Population densities recovered from beads ranged from 10^6^ to 10^9^ CFU/mL depending on species and time point. To estimate the number of generations per transfer cycle, we used the formula g = log2(N_final_/N_initial_), where N_final_ represents the population before transfer and N_initial_ represents cells on the transferred bead. Assuming a conservative starting population of ∼10^7^ cells per bead and a final population of ∼2×10^9^ cells this yields approximately 7-8 generations per cycle. With 56 transfers over 168 days, populations accumulated roughly 390-450 generations over the 24-week experiment.

**Fig. 1.**
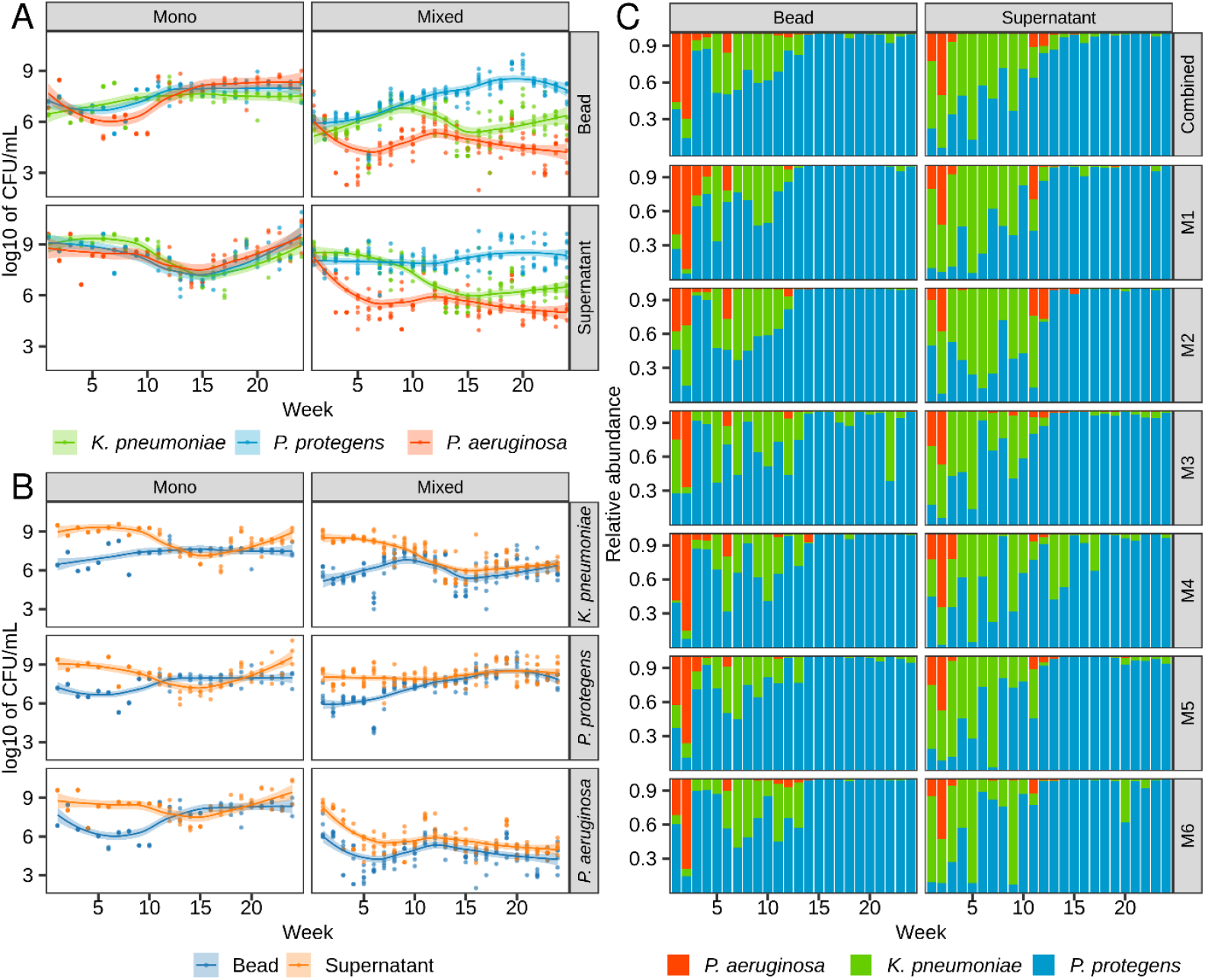
Dynamics of mono and mixed species biofilm communities during LTEE. (A, B) Temporal changes in cell counts (Log_10_ CFU/mL) of *K. pneumoniae, P. protegens* and *P. aeruginosa* recovered from biofilm beads and planktonic supernatants in mono and mixed species communities. Individual points represent absolute CFU counts, while solid lines and shaded regions depict the smoothed trajectory (loess function with 95% confidence interval). (C) Relative abundance of the three species in mixed species communities across six replicate lines (M1–M6). The first facet (top left) shows the average relative abundance across all replicates, while subsequent facets display individual community replicates.

In mono-species communities, all three species exhibited qualitatively similar population trajectories over time (Fig. 1A–B). During the early phase of evolution, planktonic populations consistently exceeded bead-associated populations by approximately one order of magnitude, indicating rapid regrowth in the supernatant following each transfer. Over successive transfers, bead-associated populations increased steadily, reflecting enhanced surface colonization and persistence. By approximately week 10–12 (corresponding to ∼100– 200 generations), cell numbers recovered from beads and supernatants converged and remained comparable through week 24. This convergence was observed across replicate lines and indicates a sustained evolutionary shift toward increased biofilm association. Importantly, these temporal dynamics are not captured by time-averaged CFU values alone but are evident from the longitudinal trajectories shown in Fig. 1.

In contrast, mixed-species communities displayed strongly divergent and time-dependent population dynamics among species (Fig. 1A–C). During the initial phase of the experiment (weeks 1–2; <50 generations), *P. aeruginosa* frequently dominated the bead-associated fraction, accounting for more than 50% of attached cells in several replicates (Fig. 1C). This early advantage was transient. From approximately week 3 onward, *P. protegens* increased sharply in bead-associated abundance and rapidly became the dominant biofilm resident. This dominance persisted for the remainder of the experiment (>200 generations), with *P. protegens* consistently comprising the majority of attached cells across all six independent mixed-community replicates (M1–M6). In contrast, *K. pneumoniae* and *P. aeruginosa* persisted at substantially lower and more variable densities within the biofilm.

Population dynamics in the supernatant phase of mixed-species communities followed a distinct temporal pattern. *K. pneumoniae* initially dominated the planktonic fraction during the early transfers, consistent with rapid post-bottleneck regrowth. However, this dominance declined over time, and from approximately week 9–10 onward *P. protegens* also became numerically dominant in the supernatant, frequently exceeding 80% relative abundance. The parallel rise of *P. protegens* in both bead-associated and planktonic fractions indicates that its evolutionary success was not restricted to surface attachment alone, but reflected a broader competitive advantage within the multispecies context.

Together, these results demonstrate that while single-species populations converge toward similar biofilm–planktonic equilibria over a few hundred generations, multispecies interactions fundamentally reshape evolutionary trajectories. Early transient dominance by one species can give way to stable long-term numerical dominance by another, emphasizing the importance of resolving population dynamics through time rather than relying on averaged abundance measures.

### Species interactions modulate the evolutionary enhancement of biofilm production

To investigate phenotypic changes during the LTEE, we profiled 648 single isolates collected at four time points (Weeks 1, 8, 15 and 24; Supplementary Table 1) using various assays as described in the Materials and Methods. These time points were selected to capture functional variation across the initial, intermediate, and final stages of the experiment. As expected, isolates from both mono and mixed species communities of *K. pneumoniae, P. protegens* and *P. aeruginosa* exhibited increased biofilm formation capacity over time (Fig. 2). At week 1, the biofilm formation ability of all three species was comparable to their respective ancestral strains, with no significant differences observed between isolates from mono and mixed species communities.

**Fig. 2.**
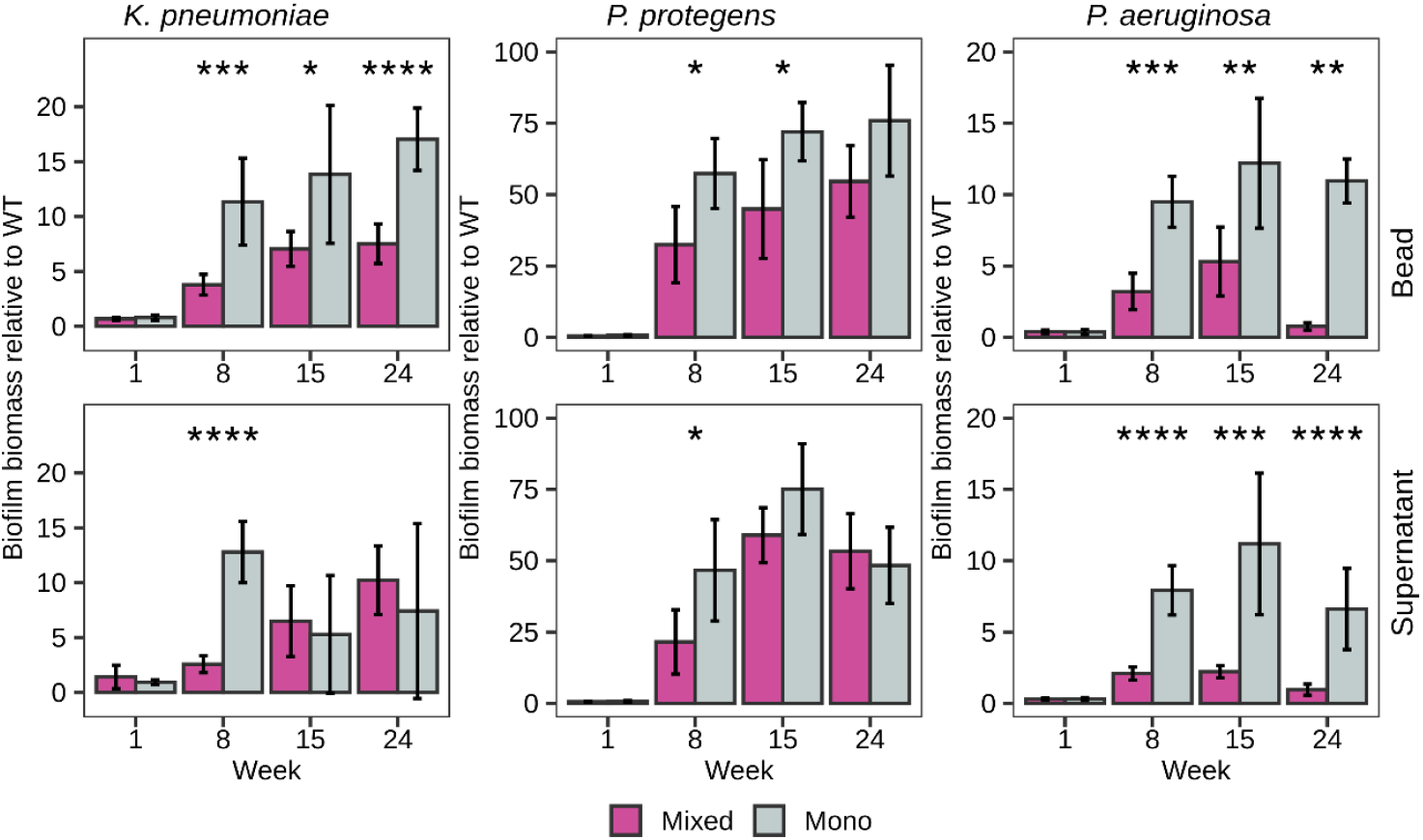
Biofilm production is constrained in mixed-species communities. The relative biofilm formation capacity compared to the respective ancestral strain are shown for the three species. Biofilm biomass was quantified using crystal violet staining assay. Data are presented as the mean of three technical replicates of respective isolates from mono and mixed communities as described in Supplementary Table 1. The error bar represents 95% CI. Statistical significance was determined using Kruskal-Wallis test between mono and mixed isolates and indicated as *, P < 0.05; **, P < 0.01; ***, P < 0.001 and ****, P < 0.0001.

In weeks 8, 15 and 24, mono isolates from beads demonstrated significantly enhanced biofilm formation compared to their mixed counterparts. For *K. pneumoniae*, mono-species and mixed-species bead isolates produced ∼17-fold and ∼6-fold increased biofilm relative to the ancestor, respectively. Among the three species, *P. protegens* exhibited the greatest capacity for biofilm production. When evolved alone on beads, *P. protegens* displayed average increases of ∼57-, ∼76- and ∼92-fold at weeks 8, 15 and 24, respectively. In contrast, when evolved in mixed communities, isolates showed ∼36-, ∼53-, and ∼57-fold increases, which proved statistically different (*P* < 0.05). Similarly, *P. aeruginosa* mono and mixed isolates showed, on average, ∼11- and ∼4-fold increases in biofilm formation, respectively. These differences were statistically significant at weeks 8 and 24 (P < 0.05).

Overall, the biofilm assays indicated that evolution in mixed communities constrained biofilm production relative to evolution in monoculture, although both sets increased significantly. *P. protegens* evolved to become the strongest biofilm former among the three species and this may have accounted for its increased abundance during long-term co-evolution.

### Mixed-species interactions constrain phenotypic diversification during biofilm evolution

To investigate whether evolution in the bead biofilm model influences phenotypes beyond biofilm formation, we performed motility, protease, and pyoverdine production assays on the two *Pseudomonas* species (Supplementary Fig. S1). *P. aeruginosa* was additionally assessed for pyocyanin production. Overall, both *P. aeruginosa* and *P. protegens* exhibited reduced motility relative to their ancestral strains during LTEE. However, no consistent differences in motility emerged between mono and mixed isolates.

Protease activity in mixed isolates remained comparable to ancestral strains, whereas bead-associated isolates from monocultures frequently displayed elevated protease production. Specifically, *P. aeruginosa* bead-associated isolates from monocultures produced significantly more protease than mixed isolates at week 24, while *P. protegens* bead-associated isolates from monocultures exceeded mixed isolates at weeks 15 and 24. Pyoverdine production remained unchanged in *P. aeruginosa* but increased over time in *P. protegens*, with mono isolates producing more than their mixed counterparts. This enhanced siderophore production likely reflects stronger iron acquisition strategies by *P. protegens* that may have contributed to its relative increase and the decline in *P. aeruginosa* abundance. In contrast, pyocyanin production in *P. aeruginosa* steadily increased over time, with mono isolates consistently surpassing mixed isolates. The parallel increase across both community contexts suggests that elevated pyocyanin production reflects a general adaptive response to the bead biofilm environment, potentially linked to metabolic activity or redox homeostasis, rather than a community-specific antagonistic strategy.

To visualize overall patterns of phenotypic variation, we performed principal component analysis (PCA) using all measured traits. Across all three species, bead-associated populations occupied a broader phenotypic space than planktonic populations, indicating greater phenotypic heterogeneity among biofilm-associated isolates (Fig. 3). Similarly, isolates evolved in monoculture showed much greater dispersion in PCA space than those from mixed-species communities, consistent with mixed-species interactions constraining phenotypic diversification. This contrast was most pronounced in *P. aeruginosa*, followed by *P. protegens* and *K. pneumoniae*.

**Fig. 3.**
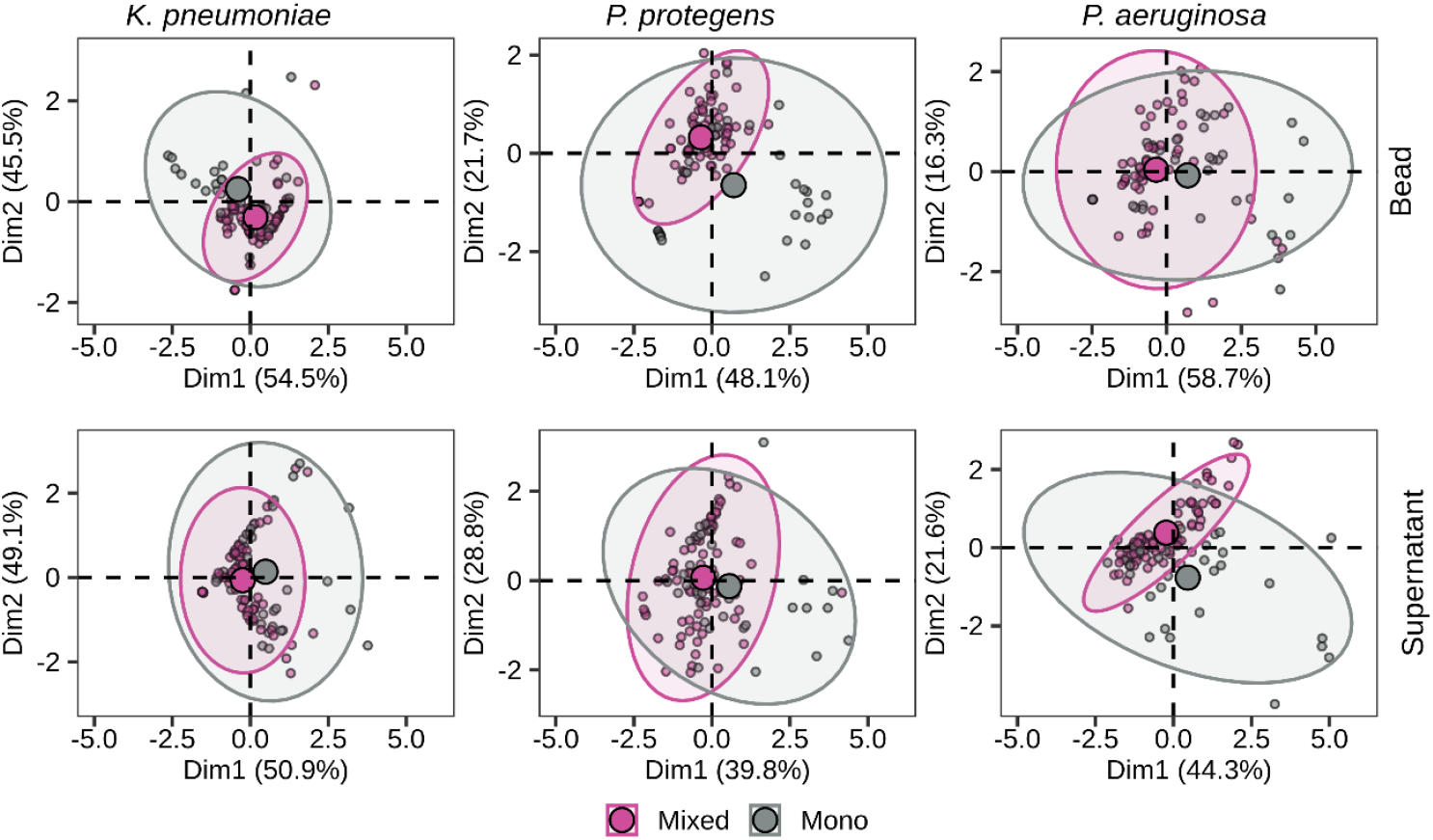
Principal component analysis (PCA) plot showing the separation of isolates from mono and mixed species communities based on phenotypic assay results. The plots display Principal Component 1 (PC1) on the x-axis and Principal Component 2 (PC2) on the y-axis and the variance are indicated in the brackets. (A) The data points are colored according to their group (mono vs. mixed). Ellipses represent the 95% confidence interval for each group. The mean point for each of the group are represented with bigger circle.

These results demonstrate that biofilm-associated evolution fosters greater phenotypic heterogeneity than planktonic growth, whereas mixed-species interactions constrain diversification. Enhanced iron acquisition traits in *P. protegens* and toxin production in *P. aeruginosa* highlight distinct adaptive strategies that likely drive shifts in community composition during LTEE.

### Genetic signatures of adaptation reveal species-specific evolutionary trajectories

To identify genetic changes underlying biofilm adaptation during the LTEE, we sequenced the genomes of the same 648 single isolates previously phenotyped at weeks 1, 8, 15, and 24. Given that mutations persisting across multiple timepoints likely represent adaptive changes under sustained selection, we focused our analysis on genes showing repeated mutations from week 8 through week 24. We also examined genes uniquely mutated in mono versus mixed communities, as these may reveal condition-specific adaptations. In addition, we performed whole-population sequencing to an average of 2400× coverage at these time points.

We identified 356, 617, and 691 mutations affecting 42, 38, and 41 genes in single isolates of *K. pneumoniae, P. protegens*, and *P. aeruginosa*, respectively. The distribution of mutations per isolate varied across species: *K. pneumoniae* isolates carried an average of 1.7 mutations (range: 0–6), *P. protegens* isolates averaged 2.3 mutations (range: 0–8), and *P. aeruginosa* isolates averaged 2.1 mutations (range: 0–7). In *K. pneumoniae*, the average number of mutations increased over time in mixed populations compared to mono isolates across both bead and supernatant conditions (Fig. 4A). By contrast, *P. protegens* and *P. aeruginosa* showed no substantial differences in mutation numbers between mono and mixed isolates.

**Fig. 4.**
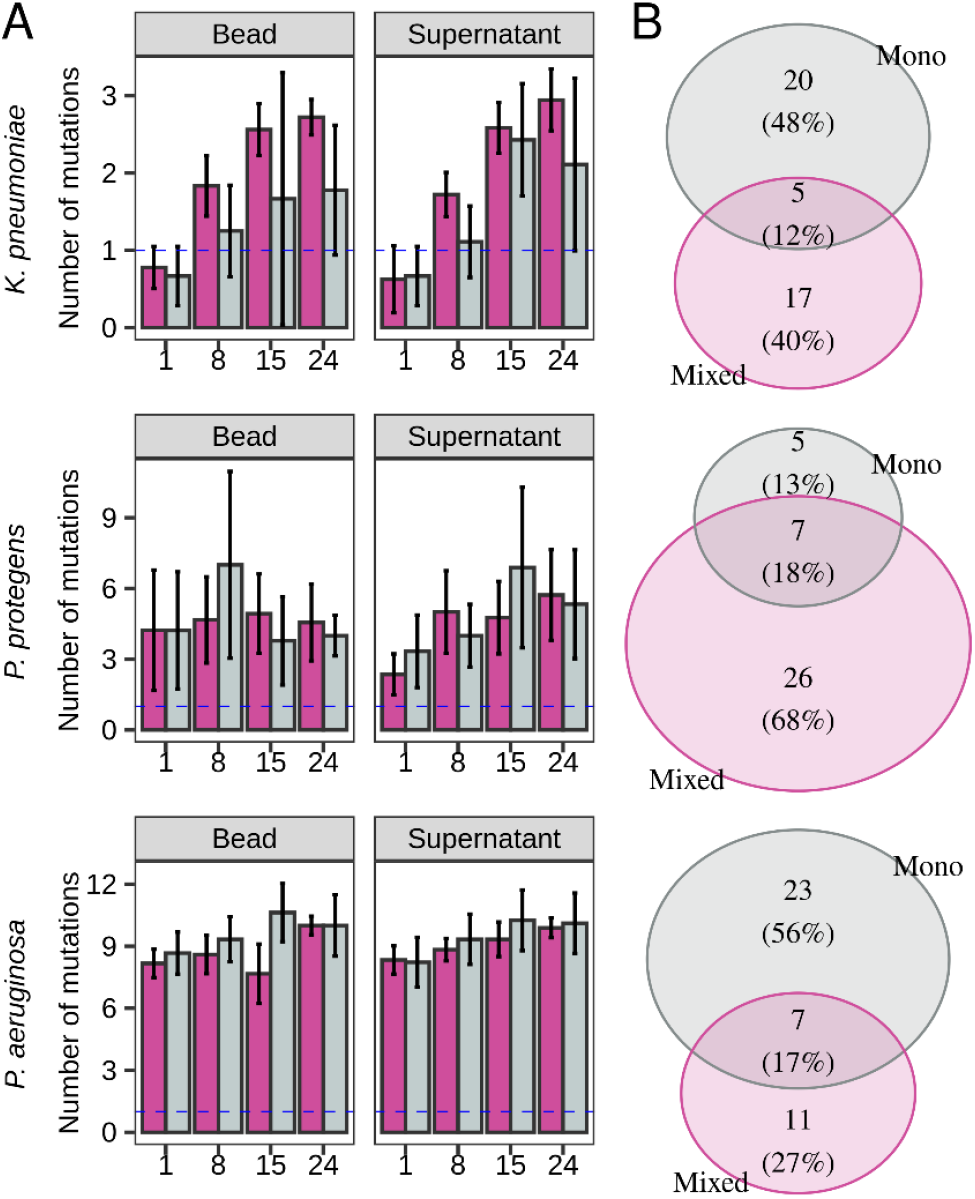
The number of mutations shared and unique genes in isolates from mono and mixed species communities. (A) The number of mutations in isolates from mono and mixed species communities across 4 time points. Data are presented as the mean and the error bar represents 95% CI. (B) The number of shared and unique genes between mono and mixed species isolates.

In *P. aeruginosa*, more genes were mutated in isolates from mono-species populations than from multispecies communities. Specifically, 56% (n = 23) of mutated genes were unique to mono isolates, 27% (n = 11) were unique to mixed isolates, and 17% (n = 7) were shared (Fig. 4B). Conversely, in *P. protegens*, fewer genes were mutated in mono-species isolates than in mixed-species isolates: 13% (n = 5) were unique to mono isolates, 68% (n = 26) were unique to mixed isolates, and 18% (n = 7) were shared. Unlike the *Pseudomonas* species, *K. pneumoniae* showed comparable proportions of unique mutations in mono (48%, n = 20) and mixed (40%, n = 17) isolates, with 12% (n = 5) of mutated genes shared between them. Analysis of mutation types revealed a higher proportion of small indels over the time in mixed species isolates of *P. protegens* and *K. pneumoniae* (Supplementary Fig. S2). In *P. aeruginosa*, small indels were the predominant mutation type overall. In *P. protegens*, non-synonymous mutations, intergenic mutations, and small indels were most common, whereas in *K. pneumoniae*, non-synonymous, synonymous, and small indel mutations predominated.

Parallel evolution, indicated by mutations arising independently at multiple timepoints, was most pronounced in *P. protegens* mixed communities, with 7 genes showing repeated mutations, compared to 2–3 genes in *K. pneumoniae* and 2–6 genes in *P. aeruginosa* (Fig. 5).

**Fig. 5.**
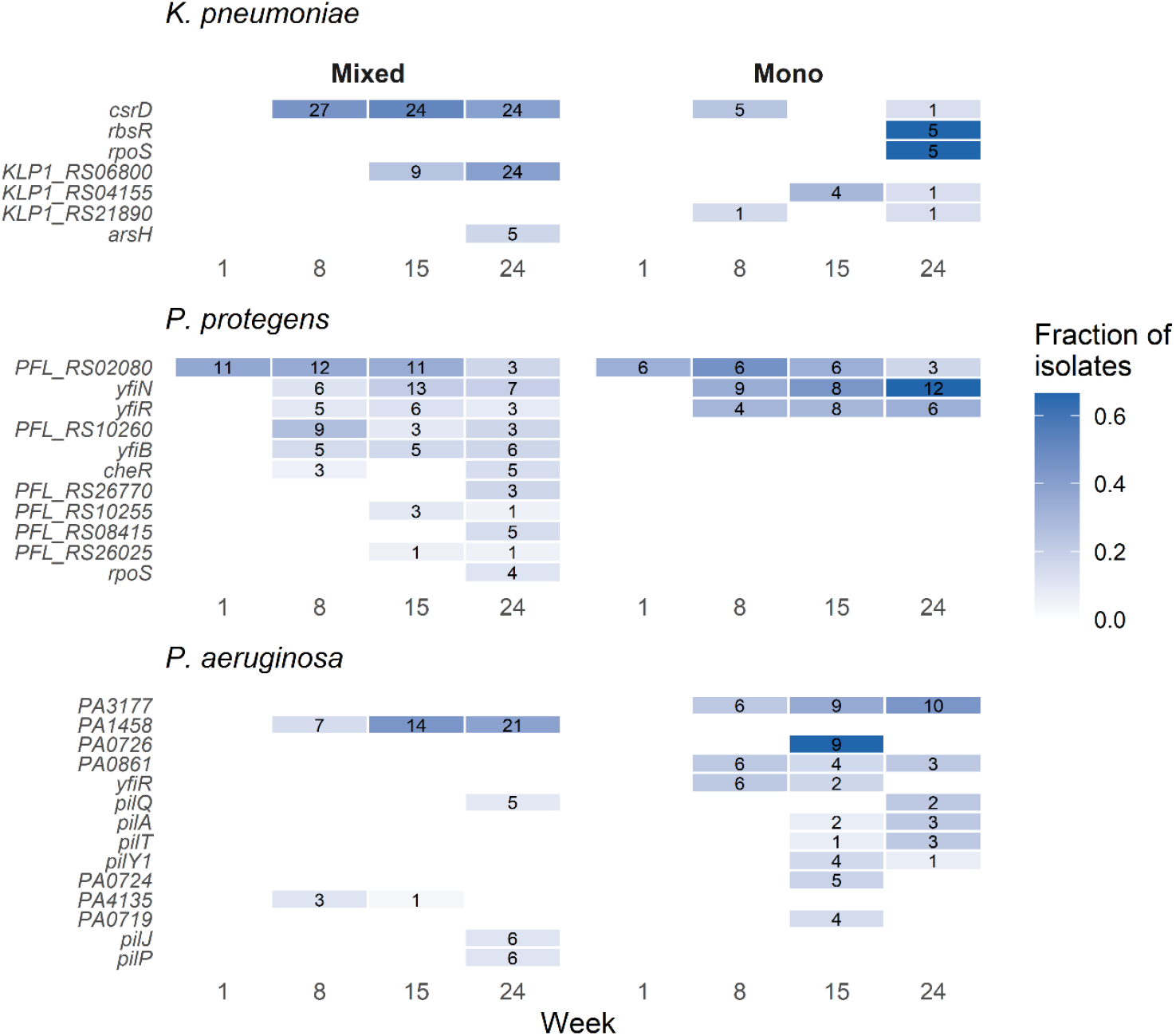
Temporal dynamics of mutations in frequently mutated genes across mono and mixed species communities. Heatmaps showing the fraction of sequenced isolates carrying non-synonymous SNPs or small indels in the most frequently mutated genes for each species. Color intensity indicates the proportion of isolates with mutations in each gene and the numbers within cells show the absolute count of mutated isolates. Only genes showing mutations in multiple independent contexts (>1 timepoint or >2 isolates) are displayed to highlight potential targets of selection. Genes are ordered by total mutation frequency across all timepoints within each species. Blank cells indicate no mutations detected.

In *K. pneumoniae*, mutations in *csrD* (carbon storage regulator/RNase E specificity factor) emerged by week 8 and persisted through week 24 (Fig. 5). Both non-synonymous mutations and small indels were detected across independent replicate populations, together accounting for 60–90% of sequenced mixed isolates from weeks 8 to 24. Notably, multiple distinct *csrD* mutations arose in parallel across different genetic backgrounds within and between replicate populations, a pattern consistent with a soft selective sweep on *csrD* function. The frequency of *csrD* mutations was comparable between bead-associated and planktonic fractions, suggesting that selection on *csrD* operates similarly across both niches. Whole-population sequencing confirmed a steady increase in *csrD* mutation frequency over time, reaching 25%, 50%, and 75% at weeks 8, 15, and 24, respectively (Supplementary Fig. S3). Although *csrD* mutations also occurred in mono isolates, they were less frequent and temporally restricted (e.g., weeks 8 and 24 in bead-associated isolates, week 8 only in supernatant isolates). An in-depth analysis identified a dominant nSNP at position 938 (A→G) in *csrD* (Supplementary Fig. S4).

In *P. protegens*, mutations were observed in the *yfiBNR* operon (*yfiB, yfiN, yfiR*) from weeks 8 to 24, with this operon showing parallel mutations across 3 timepoints in mixed communities. YfiBNR is a tripartite signaling module that controls intracellular c-di-GMP levels and small colony variant formation [22]. Notably, *yfiB* mutations occurred exclusively in mixed isolates, affecting ∼20% of bead-associated isolates, whereas *yfiN* and *yfiR* mutations were present in both mono and mixed isolates. Whole-population data mirrored these patterns, with *yfiB* mutated only in mixed populations, while *yfiN* and *yfiR* mutations were distributed across both evolutionary contexts (Supplementary Fig. S3). The most prevalent mutation among mixed *P. protegens* isolates was a nSNP at position 94 (A→C) in *yfiB*, resulting in an asparagine-to-histidine substitution (N32H) in the periplasmic domain. This residue lies within a region critical for YfiB’s interaction with YfiR, suggesting that the mutation may disrupt sequestration of YfiR and consequently elevate c-di-GMP levels to promote biofilm formation.

In *P. aeruginosa*, small indel mutations in the chemotaxis gene *cheA* were unique to mixed isolates and persisted from week 8 to week 24. At week 8, 30% of bead-associated and 10% of planktonic isolates carried *cheA* mutations; by week 15, these proportions rose to 100% and 60%, respectively, before declining slightly at week 24. Conversely, *dgcN* (PA3177), encoding a diguanylate cyclase, was mutated only in mono isolates, with frequencies increasing over time across 3 timepoints. In bead-associated mono isolates, *dgcN* mutation prevalence rose from 30% (week 8) to 60% (week 15) and 80% (week 24); a similar upward trend was observed in planktonic isolates (30%, 50%, and 60% over the same intervals). In addition to *dgcN*, mutations in PA0861 (*rbdA*), were detected in mono isolates (Supplementary Fig. S3). Unlike DgcN, RbdA functions as a phosphodiesterase that degrades c-di-GMP and promotes biofilm dispersal. The concurrent mutations in both c-di-GMP synthesis (*dgcN*) and degradation (*rbdA*) pathways exclusively in monocultures suggest that fine-tuning intracellular c-di-GMP levels is a critical adaptive strategy for *P. aeruginosa* biofilm regulation in the absence of interspecific competition. Whole-population sequencing confirmed an increasing frequency of nSNPs in *dgcN* across mono populations.

Collectively, our findings identify *csrD, yfiBNR*, and *cheA* as repeatedly selected genetic targets in *K. pneumoniae, P. protegens*, and *P. aeruginosa*, respectively. The temporal dynamics of these mutations emerging early and increasing in frequency over 24 weeks combined with evidence of parallel evolution across independent lineages, strongly support their adaptive significance in biofilm formation. Species-specific mutation patterns, particularly the enrichment of *yfiB* and *cheA* mutations exclusively in mixed communities, suggest distinct genetic routes to biofilm adaptation shaped by interspecific interactions.

### Mutations in *csrD, yfiBNR* and *cheA* drive enhanced biofilm formation

Given that mutations in *csrD, yfiBNR* and *cheA* (PA1458) were the most prevalent in mixed isolates of *K. pneumoniae, P. protegens*, and *P. aeruginosa*, respectively, we hypothesized that these genetic changes play a major role in biofilm formation. To test this, we categorized phenotypic data based on the presence or absence of mutations in these genes. Isolates carrying mutations in *csrD, yfiBNR* or *cheA* exhibited a significant increase in biofilm formation capacity compared with those lacking such mutations (Fig. 6A). Specifically, *K. pneumoniae* isolates with *csrD* mutations showed an approximately 7.5-fold increase in biofilm formation relative to non-*csrD* isolates (P < 0.05). *P. protegens* isolates with *yfiBNR* mutations displayed an approximately 3.5-fold increase (P < 0.05), and *P. aeruginosa* isolates with *cheA* mutations showed a ∼2-fold increase (P < 0.05).

**Fig. 6.**
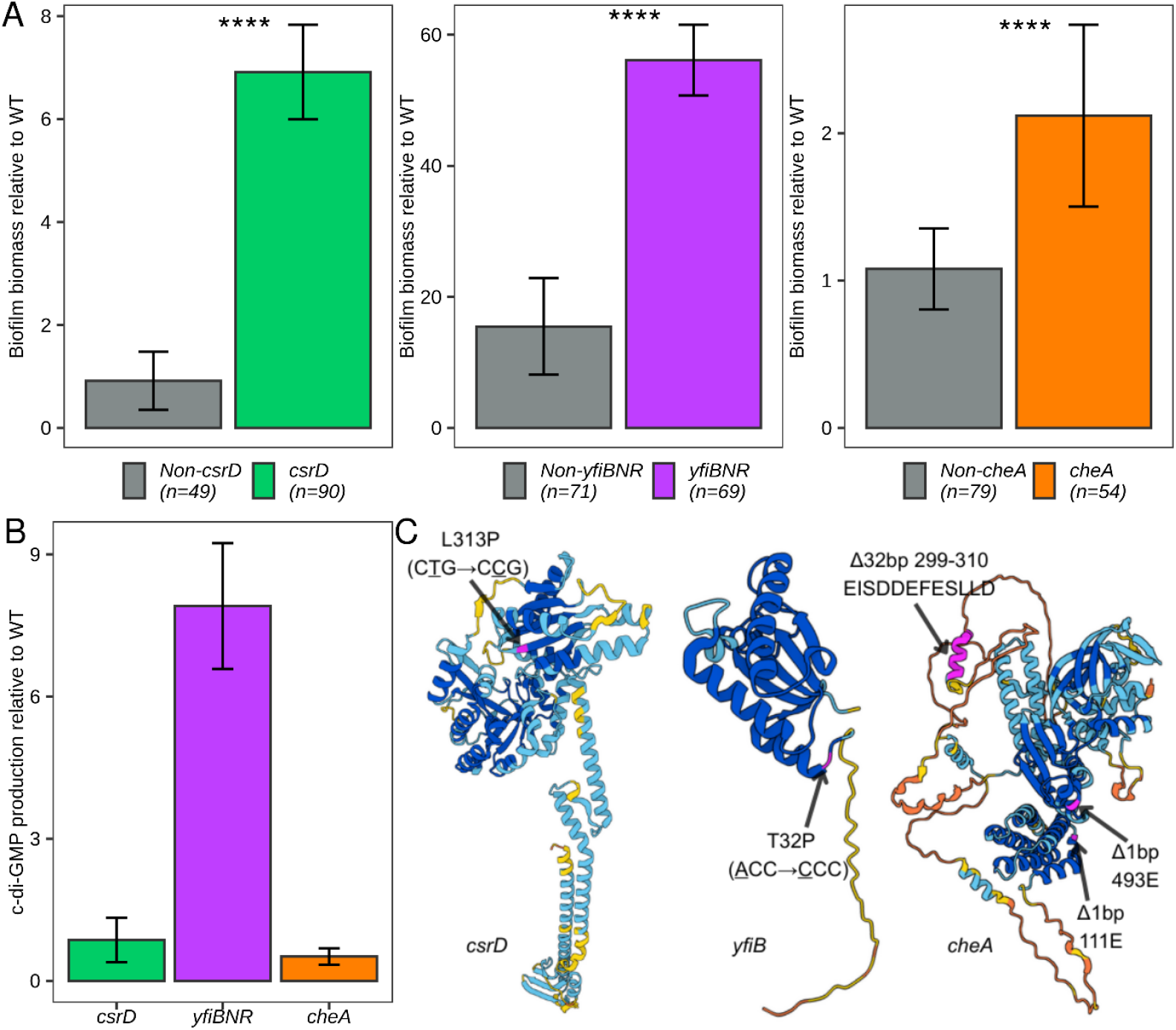
Mutations in *csrD, yfiBNR* and *cheA* lead to the enhanced biofilm formation and cyclic-di-GMP production. (A) Biofilm biomass of isolates with and without mutations in the respective key genes for each species. The number of isolates for each of the categories are shown in the legend in brackets. Biofilm biomass were measured from three technical replicates per isolate using crystal violet staining. (B) Cyclic-di-GMP production capacity of the mixed isolates of *K. pneumoniae, P. protegens* and *P. aeruginosa* with mutations in *csrD* (n=3), *yfiBNR* (n=3) and *cheA* (n=3), respectively. Cyclic-di-GMP were quantified using ELISA kit and expressed in pg/mL. The relative c-di-GMP production capacity compared to the respective wild-type strain are shown. (C) Alphafold predicted protein structural model of CsrD, YfiB and CheA highlighting the mutations of interest in pink. Data are presented as means, with error bars indicating 95% confidence intervals. Statistical significance was determined using the Kruskal–Wallis test: *, P < 0.05; **, P < 0.01; ***, P < 0.001; ****, P < 0.0001.

Since increased biofilm formation is often associated with elevated intracellular c-di-GMP, we quantified c-di-GMP levels in isolates carrying mutations in *csrD, yfiBNR*, and *cheA*. The assay revealed that *P. protegens* isolates with mutations in *yfiBNR* exhibited an approximately 8-fold increase in c-di-GMP levels compared to the wild-type strain (Fig. 6B). In contrast, *K. pneumoniae* and *P. aeruginosa* isolates harboring mutations in *csrD* and *cheA*, respectively, showed no substantial change in c-di-GMP production relative to their wild-type counterparts.

Structural analysis of the mutated proteins provided further insights. The dominant nSNP identified in the *yfiB* gene resulted in an amino acid substitution (T32P, ACC → CCC) with a high confidence prediction (pLDDT > 90; Fig. 6C). This mutation is located within the N-terminal region of YfiB, which has previously been implicated in interactions with YfiR. Earlier studies have shown that single amino acid substitutions in the N-terminal domain of YfiB can promote conformational changes that facilitate sequestration of YfiR into a stable complex [23, 24]. Such sequestration relieves YfiR-mediated repression of YfiN, ultimately leading to increased c-di-GMP production.

### Mutations in *yfiBNR* are the strongest driver in shaping long-term co-evolutionary dynamics

We further hypothesized that these mutations also influence the relative abundance of the three species within mixed communities, driving the species dynamics observed during the LTEE. To investigate this, we selected mixed isolates of *K. pneumoniae, P. protegens* and *P. aeruginosa* carrying the most prevalent mutation types (presented in Supplementary Fig. S4) in *csrD, yfiBNR* and *cheA*, respectively and performed competition assays under LTEE-like conditions. Strikingly, mutations in *yfiBNR* were associated with a dominance of *P. protegens* in the biofilm fraction, closely mirroring LTEE community composition (Fig. 7A). In contrast, competitions with wild-type strains or week-1 isolates resulted in high *P. aeruginosa* abundance in the bead fraction and high *K. pneumoniae* abundance in the supernatant. When *yfiBNR*-mutated isolates were present, *P. protegens* constituted ∼60–80% of the bead-associated population, whereas their absence led to increased bead-associated abundance of *K. pneumoniae*, followed by *P. aeruginosa*. In the supernatant fraction, *K. pneumoniae* dominated regardless of the specific mutant combination.

**Fig. 7.**
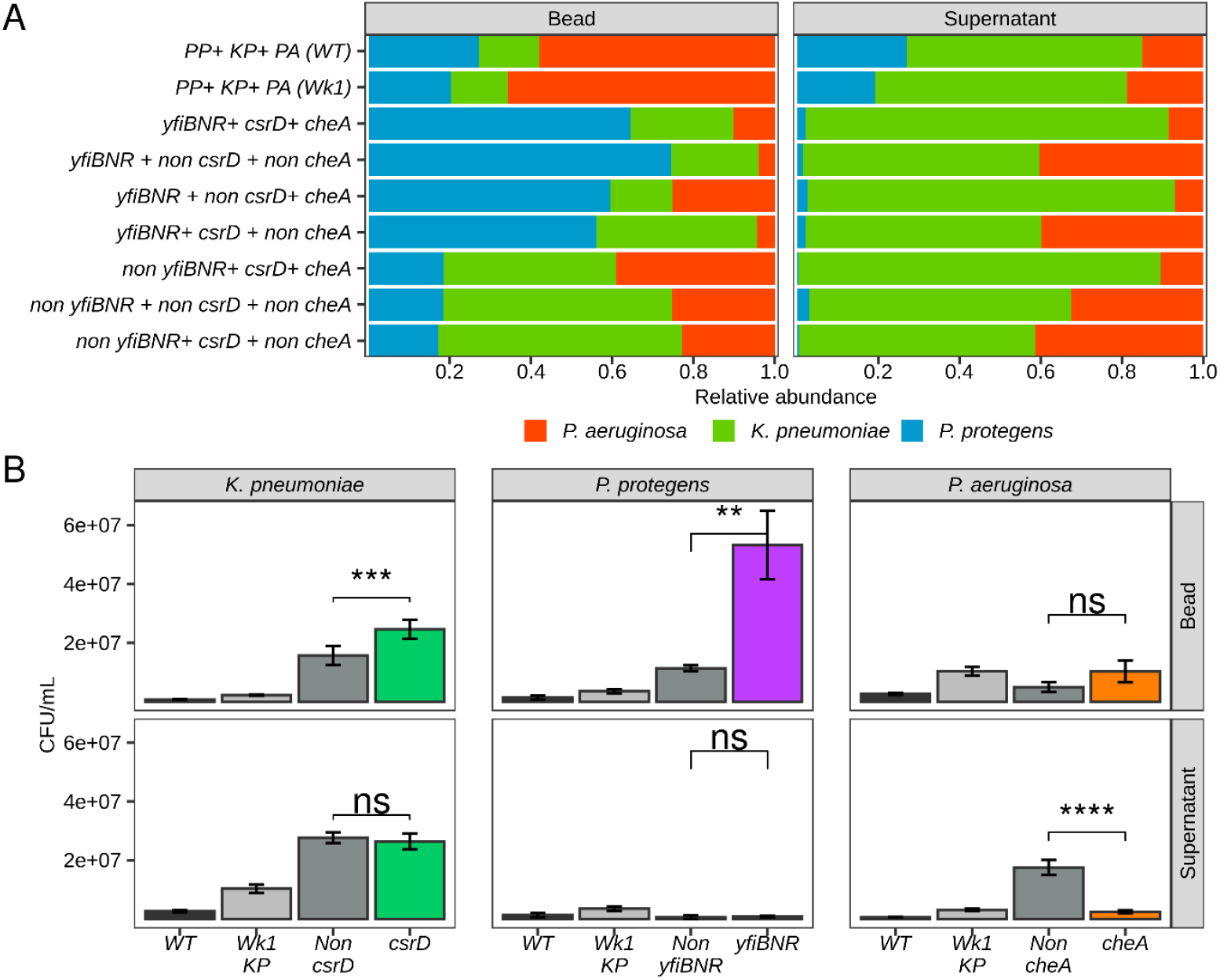
Mutations in *csrD, yfiBNR*, and *cheA* act as key drivers of the observed community dynamics during LTEE. (A) Relative abundance of mixed isolates of *K. pneumoniae, P. protegens* and *P. aeruginosa* with and without mutations in *csrD, yfiBNR* and *cheA*, respectively. (B) Differences in cell counts during competition assays between mixed isolates of the three species. Data are presented as means, with error bars indicating 95% confidence intervals. Statistical significance was determined using the Kruskal–Wallis test: *, P < 0.05; **, P < 0.01; ***, P < 0.001; ****, P < 0.0001.

Quantitative cell counts revealed that *K. pneumoniae* isolates with *csrD* mutations had significantly higher bead-associated abundance compared to non-*csrD* isolates (Fig. 7B). Similarly, *yfiBNR*-mutated *P. protegens* displayed significantly higher bead attached cells relative to non-mutated counterparts. Although *cheA*-mutated *P. aeruginosa* had slightly higher bead-attached counts than non-mutated isolates, the difference was not statistically significant. In contrast, non-*cheA* isolates had significantly greater planktonic abundance in the compared to *cheA*-mutated isolates.

Collectively, these results reveal distinct roles for *csrD, yfiBNR* and *cheA* mutations in structuring biofilm and planktonic populations within multispecies communities. While all three mutations enhance biofilm formation, *yfiBNR* mutations in *P. protegens* exerted the strongest effect, driving competitive dominance and shaping community composition in a manner consistent with patterns observed during LTEE. This suggests that specific genetic changes can act as keystone drivers of both phenotypic adaptation and long-term ecological dynamics in mixed-species biofilms.

## Discussion

Our LTEE of mono- and mixed-species biofilm communities reveals that the trajectory of microbial adaptation is profoundly shaped by species interactions. In monoculture, all three species, *K. pneumoniae, P. protegens* and *P. aeruginosa*, evolved towards increased biofilm production and phenotypic diversification via distinct genetic pathways. In contrast, mixed-species communities exhibited constrained phenotypic diversity and altered population dynamics, with *P. protegens* emerging as the dominant biofilm resident over 24 weeks. These findings demonstrate that mixed-species interactions can both limit the breadth of adaptive evolution and concentrate selection pressure on specific genetic loci that influence community composition and stability, particularly those governing biofilm formation and competitive interactions.

### Species interactions reshape adaptive landscapes

In mono species communities, biofilm biomass increased steadily over time, consistent with selection for traits that enhance attachment and persistence in structured habitats [25, 26]. By contrast, in mixed-species communities, the extent of increased biofilm production was limited in all species, suggesting that interspecific competition or antagonism imposes constraints on this trait. The ecological consequences of these altered evolutionary paths were evident in the community dynamics. In co-evolution, *P. protegens* progressively displaced both *K. pneumoniae* and *P. aeruginosa* from bead-associated biofilms, despite not dominating in monoculture. This shift coincided with the emergence of mutations in the *yfiBNR* operon, an established central regulator of cyclic-di-GMP signaling and biofilm formation in *Pseudomonas* sp. [22, 27]. The dominance of *P. protegens* in both beads and supernatants from week 10 onwards suggests that biofilm-associated advantages, possibly coupled with enhanced iron acquisition, conferred a competitive edge in the mixed-species context. This outcome is consistent with the bead transfer model’s inherent selection for surface-associated traits.

### Keystone mutations drive phenotypic and ecological outcomes

Whole-genome sequencing revealed that specific mutations in *csrD* (*K. pneumoniae*), *yfiBNR* (*P. protegens*) and *cheA* (*P. aeruginosa*) arose early and persisted across the LTEE. Competition assays confirmed that these mutations enhanced biofilm formation, with *yfiBNR* variants in *P. protegens* exerting the largest effect on competitive dominance. The *csrD* mutations likely impact the turnover of regulatory RNAs controlling glycogen storage and biofilm-related pathways, while *cheA* indels in *P. aeruginosa* may alter chemotaxis responses, shifting the balance between surface colonization and dispersal [28].

Mutations targeting global regulatory networks and cyclic-di-GMP signaling pathways are a recurrent outcome of biofilm-focused experimental evolution [29, 30], and our findings align closely with this pattern. Importantly, however, the ecological effects of such mutations were strongly context dependent. For example, although *cheA* mutations were associated with increased biofilm formation, these variants did not confer dominance in the bead fraction when competing against *yfiBNR*-mutated *P. protegens*. This highlights that the fitness effect of a given mutation cannot be inferred from monoculture performance alone and depends critically on the identity and evolutionary state of competing species. More broadly, our results support the growing recognition that adaptive value in microbial evolution is shaped by ecological context, where competitive hierarchies and spatial structure determine which mutations translate into long-term persistence [31].

### Mixed-species biofilms constrain diversification

Phenotypic profiling and PCA demonstrated that monoculture evolution yielded greater trait diversity than co-evolution, particularly in bead-associated populations. This aligns with theory predicting that competitive interactions in fluctuating environments can favor a narrower set of successful strategies when strong competitors emerge, thereby reducing diversification [32, 33]. The observed patterns in our system including the reduced motility and elevated iron acquisition in *P. protegens* alongside *P. aeruginosa* investment in secondary metabolite production (pyocyanin) are consistent with niche specialization. However, further experiments would be needed to definitively establish whether these represent adaptive trade-offs or simply the most accessible evolutionary paths under the selection regime imposed by the bead transfer protocol. The constraint on phenotypic diversity in mixed-species populations likely arises from both ecological filtering, where only certain phenotypes persist under competitive pressure, and evolutionary pressure, where the fitness landscape is narrowed by persistent interspecific interactions. Importantly, such constraints do not preclude high community productivity or stability, as evidenced by the stable bead/supernatant ratios from mid-experiment onwards. Instead, they reflect an evolutionary trade-off between diversity and competitive exclusion.

### Implications for natural and applied systems

Our findings have direct implications for understanding microbial community assembly and stability in natural biofilms and engineered systems. In multispecies contexts, keystone mutations such as those identified in *yfiBNR* can rapidly and persistently reshape community structure, potentially overriding the effects of initial community composition. This echoes observations from clinical and environmental biofilms, where a small subset of genotypes can dominate over extended periods [34, 35].

From an applied perspective, our results suggest that interventions targeting specific regulatory pathways could selectively disrupt dominant biofilm formers in polymicrobial infections or industrial biofilms. Conversely, promoting beneficial keystone genotypes may stabilize desired community functions in bioreactors or plant microbiomes.

Overall, this work underscores that microbial evolution in structured, multispecies environments cannot be predicted from single-species trajectories alone. Instead, species interactions both constrain and direct adaptation, channeling evolution towards a limited set of successful ecological strategies driven by a small number of high-impact mutations. Understanding these evolutionary “shortcuts” will be essential for forecasting and manipulating biofilm community outcomes in both natural and engineered settings.

## Acknowledgements

We are thankful to other members of McDougald lab for their valuable suggestions. This work was supported by Australian Research Council Discovery Project DP230101760 awarded to Diane McDougald. We are thankful to UTS high performance computing facility HPC for data analysis.

## Author contributions

MMH, JT, VSC, SAR and DM designed the study. MMH, SH, JT, PN, and GEV performed the experiments and analyzed and interpreted the data. MMH took the lead to write the manuscript and bioinformatic analyses. VSC, SAR and DM provided funding. All authors reviewed and provided critical feedback of the manuscript.

## References

1. McDougald D, Rice SA, Barraud N, Steinberg PD, Kjelleberg S. Should we stay or should we go: mechanisms and ecological consequences for biofilm dispersal. Nature Reviews Microbiology. 2012;10(1):39–50.

2. Poltak SR, Cooper VS. Ecological succession in long-term experimentally evolved biofilms produces synergistic communities. The ISME journal. 2011;5(3):369–78.

3. Hall-Stoodley L, Costerton JW, Stoodley P. Bacterial biofilms: from the natural environment to infectious diseases. Nature reviews microbiology. 2004;2(2):95–108.

4. Flemming H-C, Wingender J, Szewzyk U, Steinberg P, Rice SA, Kjelleberg S. Biofilms: an emergent form of bacterial life. Nature Reviews Microbiology. 2016;14(9):563–75.

5. Xavier JB, Foster KR. Cooperation and conflict in microbial biofilms. Proceedings of the National Academy of Sciences. 2007;104(3):876–81.

6. Lee KWK, Periasamy S, Mukherjee M, Xie C, Kjelleberg S, Rice SA. Biofilm development and enhanced stress resistance of a model, mixed-species community biofilm. The ISME journal. 2014;8(4):894–907.

7. Mitri S, Richard Foster K. The genotypic view of social interactions in microbial communities. Annual review of genetics. 2013;47(1):247–73.

8. Booth SC, Rice SA. Influence of interspecies interactions on the spatial organization of dual species bacterial communities. Biofilm. 2020;2:100035.

9. Lenski RE, Rose MR, Simpson SC, Tadler SC. Long-term experimental evolution in Escherichia coli. I. Adaptation and divergence during 2,000 generations. The American Naturalist. 1991;138(6):1315–41.

10. Lenski RE. Experimental evolution and the dynamics of adaptation and genome evolution in microbial populations. The ISME journal. 2017;11(10):2181–94.

11. Oliveira NM, Niehus R, Foster KR. Evolutionary limits to cooperation in microbial communities. Proceedings of the National Academy of Sciences. 2014;111(50):17941–6.

12. Mitri S, Clarke E, Foster KR. Resource limitation drives spatial organization in microbial groups. The ISME journal. 2016;10(6):1471–82.

13. Madsen JS, Røder HL, Russel J, Sørensen H, Burmølle M, Sørensen SJ. Coexistence facilitates interspecific biofilm formation in complex microbial communities. Environmental microbiology. 2016;18(8):2565–74.

14. Palleroni NJ: The pseudomonas story. In., vol. 12: Wiley Online Library; 2010: 1377–83.

15. Anand V, Sankar P, Vasan R. Isolation and characterization of bacteria from the gut Of Bombyx Mori that degrade cellulose, xylan, pectin and starch and their impact on digestion. J of Insect Science. 2009;10(107):1–20.

16. Diggle SP, Griffin AS, Campbell GS, West SA. Cooperation and conflict in quorum-sensing bacterial populations. Nature. 2007;450(7168):411–4.

17. Elias S, Banin E. Multi-species biofilms: living with friendly neighbors. FEMS microbiology reviews. 2012;36(5):990–1004.

18. Subramoni S, Muzaki MZBM, Booth SC, Kjelleberg S, Rice SA. N-acyl homoserine lactone-mediated quorum sensing regulates species interactions in multispecies biofilm communities. Frontiers in Cellular and Infection Microbiology. 2021;11:646991.

19. Gaio D, Anantanawat K, To J, Liu M, Monahan L, Darling AE. Hackflex: low-cost, high-throughput, Illumina Nextera Flex library construction. Microbial Genomics. 2022;8(1):000744.

20. Deatherage DE, Barrick JE. Identification of mutations in laboratory-evolved microbes from next-generation sequencing data using breseq. Methods Mol Biol. 2014:165–88.

21. Varadi M, Bertoni D, Magana P, Paramval U, Pidruchna I, Radhakrishnan M, et al. AlphaFold Protein Structure Database in 2024: providing structure coverage for over 214 million protein sequences. Nucleic acids research. 2024;52(D1):D368–D75.

22. Malone JG, Jaeger T, Spangler C, Ritz D, Spang A, Arrieumerlou C, et al. YfiBNR mediates cyclic di-GMP dependent small colony variant formation and persistence in Pseudomonas aeruginosa. PLoS pathogens. 2010;6(3):e1000804.

23. Xu M, Yang X, Yang X-A, Zhou L, Liu T-Z, Fan Z, et al. Structural insights into the regulatory mechanism of the Pseudomonas aeruginosa YfiBNR system. Protein & Cell. 2016;7(6):403–16.

24. Li S, Li T, Xu Y, Zhang Q, Zhang W, Che S, et al. Structural insights into YfiR sequestering by YfiB in Pseudomonas aeruginosa PAO1. Scientific reports. 2015;5(1):16915.

25. Foster KR, Bell T. Competition, not cooperation, dominates interactions among culturable microbial species. Current biology. 2012;22(19):1845–50.

26. Dragoš A, Kovács ÁT. The peculiar functions of the bacterial extracellular matrix. Trends in microbiology. 2017;25(4):257–66.

27. Merritt JH, Ha D-G, Cowles KN, Lu W, Morales DK, Rabinowitz J, et al. Specific control of Pseudomonas aeruginosa surface-associated behaviors by two c-di-GMP diguanylate cyclases. MBio. 2010;1(4):10.1128/mbio.00183-10.

28. Wadhams GH, Armitage JP. Making sense of it all: bacterial chemotaxis. Nature reviews Molecular cell biology. 2004;5(12):1024–37.

29. Traverse CC, Mayo-Smith LM, Poltak SR, Cooper VS. Tangled bank of experimentally evolved Burkholderia biofilms reflects selection during chronic infections. Proceedings of the National Academy of Sciences. 2013;110(3):E250–E9.

30. Flynn KM, Dowell G, Johnson TM, Koestler BJ, Waters CM, Cooper VS. Evolution of ecological diversity in biofilms of Pseudomonas aeruginosa by altered cyclic diguanylate signaling. Journal of bacteriology. 2016;198(19):2608–18.

31. Bailey SF, Kassen R. Spatial structure of ecological opportunity drives adaptation in a bacterium. The American Naturalist. 2012;180(2):270–83.

32. Hardin G. The competitive exclusion principle: an idea that took a century to be born has implications in ecology, economics, and genetics. science. 1960;131(3409):1292–7.

33. Hibbing ME, Fuqua C, Parsek MR, Peterson SB. Bacterial competition: surviving and thriving in the microbial jungle. Nature reviews microbiology. 2010;8(1):15–25.

34. Markussen T, Marvig RL, Gómez-Lozano M, Aanæs K, Burleigh AE, Høiby N, et al. Environmental heterogeneity drives within-host diversification and evolution of Pseudomonas aeruginosa. MBio. 2014;5(5):10.1128/mbio.01592-14.

35. Ren D, Madsen JS, Sørensen SJ, Burmølle M. High prevalence of biofilm synergy among bacterial soil isolates in cocultures indicates bacterial interspecific cooperation. The ISME journal. 2015;9(1):81–9.

