## Supplementary material for "Mixed-species interactions constrain diversification and shape biofilm evolution": Mixed_species_biofilm_manuscript_Supplementary_File_biorxiv.pdf

Running Title: Long-term evolution of a mixed species biofilms.

**Keywords:** Biofilm, LTEE, diversity, evolution, phenotypes, mutation.

**Content:**

Supplementary Table 1
Supplementary Figures S1-S4

**Supplementary Table 1: Number of isolates phenotyped across different condition.**

| Evolved/Co-evolved | Bead/Supernatant | Week | *K. pneumoniae* | *P. protegens* | *P. aeruginosa* |
|---|---|---|---|---|---|
| Co-evolved | Bead | 1 | 18 | 18 | 18 |
| Co-evolved | Bead | 8 | 18 | 18 | 18 |
| Co-evolved | Bead | 15 | 18 | 18 | 18 |
| Co-evolved | Bead | 24 | 18 | 18 | 18 |
| Co-evolved | Supernatant | 1 | 18 | 18 | 18 |
| Co-evolved | Supernatant | 8 | 18 | 18 | 18 |
| Co-evolved | Supernatant | 15 | 18 | 18 | 18 |
| Co-evolved | Supernatant | 24 | 18 | 18 | 18 |
| Evolved | Bead | 1 | 9 | 9 | 9 |
| Evolved | Bead | 8 | 9 | 9 | 9 |
| Evolved | Bead | 15 | 9 | 9 | 9 |
| Evolved | Bead | 24 | 9 | 9 | 9 |
| Evolved | Supernatant | 1 | 9 | 9 | 9 |
| Evolved | Supernatant | 8 | 9 | 9 | 9 |
| Evolved | Supernatant | 15 | 9 | 9 | 9 |
| Evolved | Supernatant | 24 | 9 | 9 | 9 |

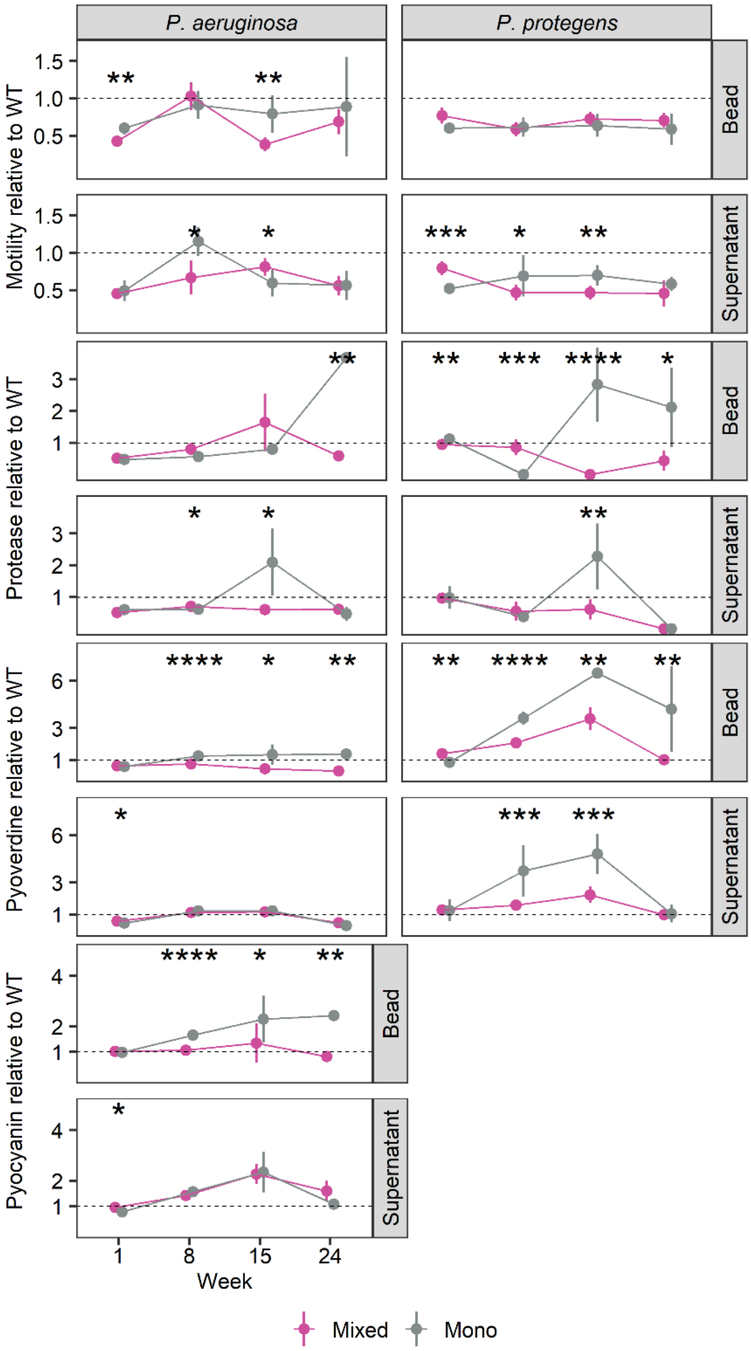


**Supplementary Fig. S1: Phenotypes of *Pseudomonas* spp.**

Data are presented as the mean of three technical replicates of respective evolved and co-evolved isolates as described in Supplementary Table 1. The error bar represents 95% CI. Statistical significance was determined using Kruskal-Wallis test between evolved and co-evolved isolates and indicated as *, P < 0.05; **, P < 0.01; ***, P < 0.001 and ****, P < 0.0001.

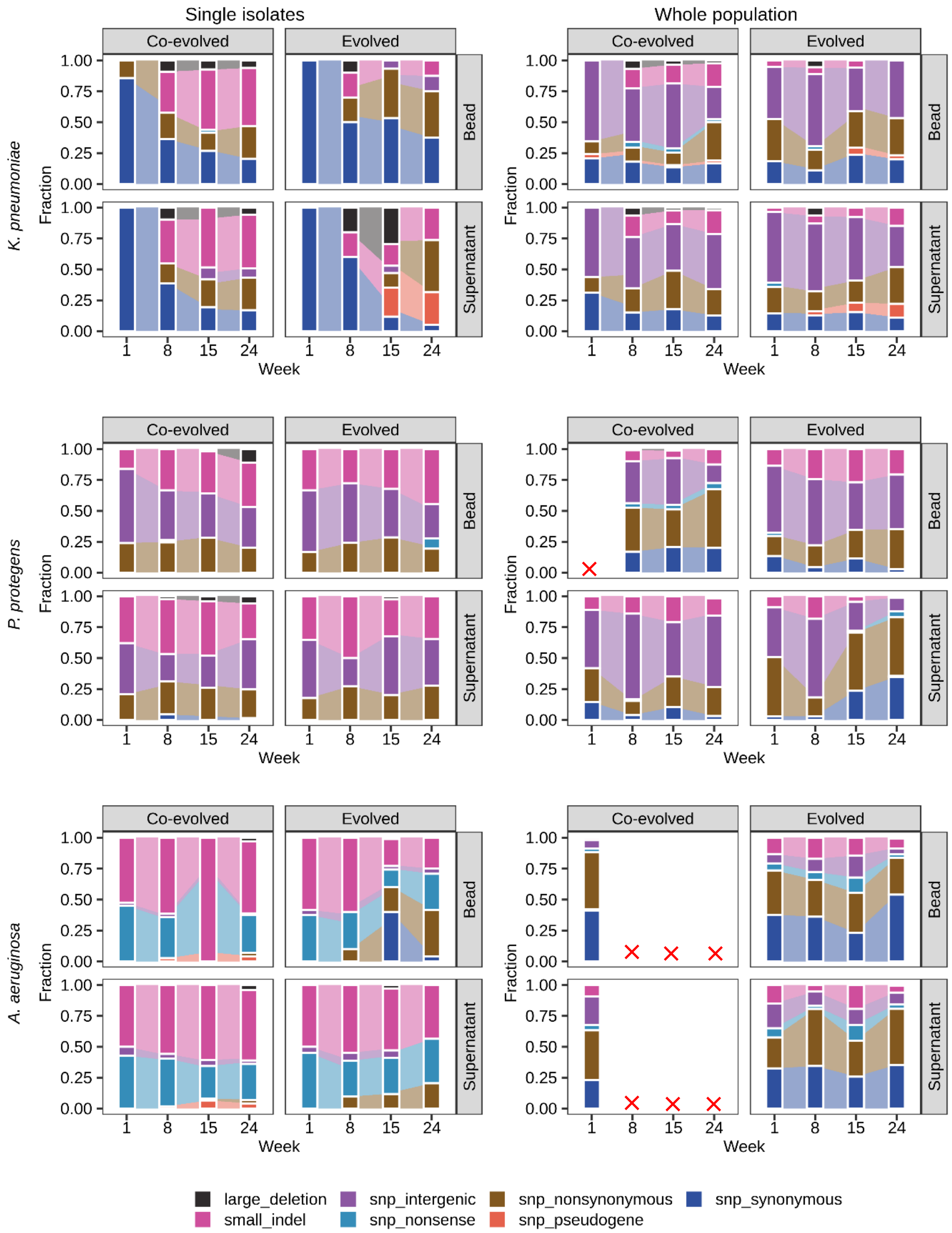


**Supplementary Fig. S2: Mutation profiles of evolved and co-evolved isolates.** This stacked bar chart displays the relative frequency of different mutation types, with each color representing a specific mutation type as defined in the legend. A red cross indicates insufficient sequencing reads for the corresponding species and week, leading to no data analysis for that point.

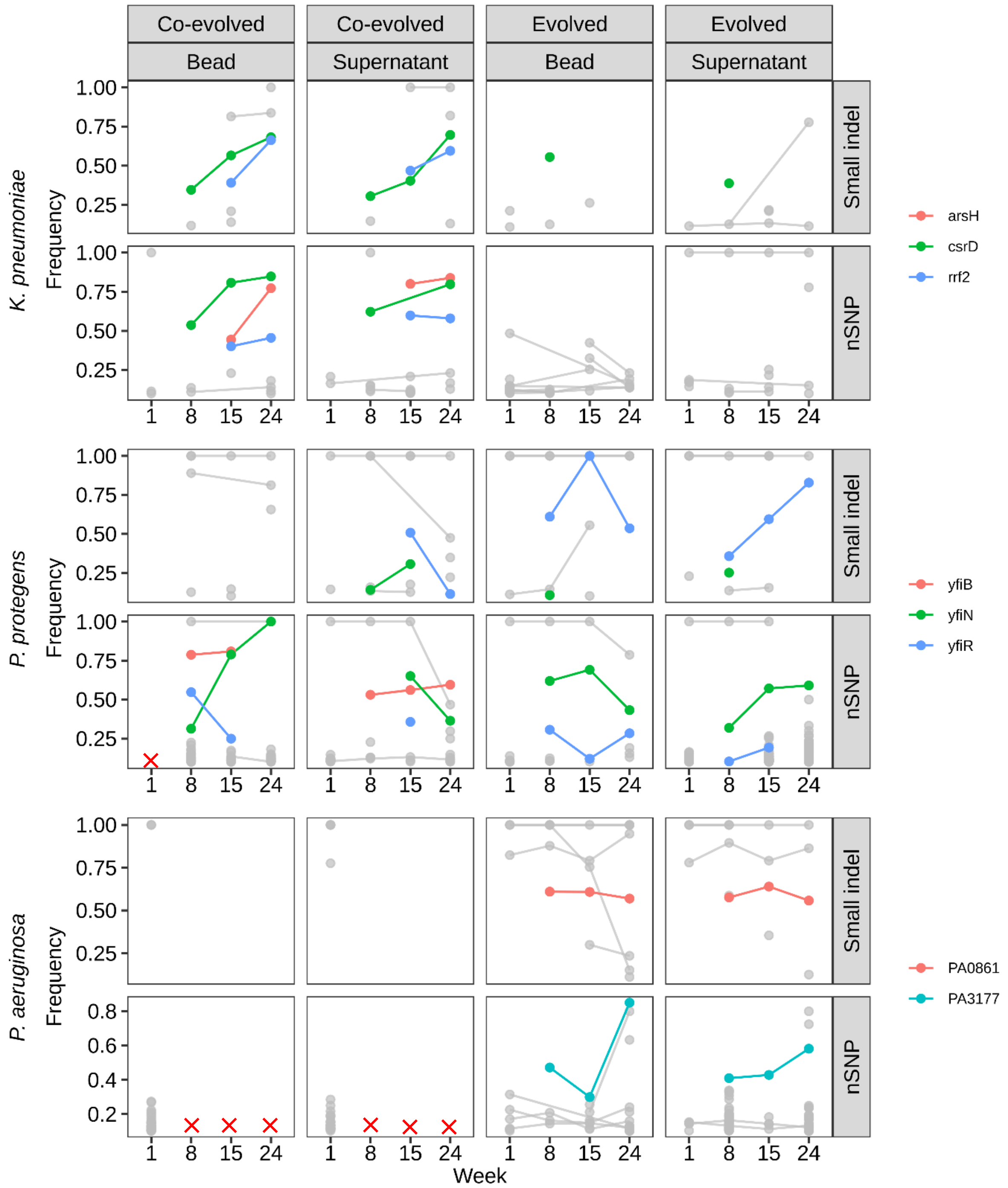


**Supplementary Fig. S3: Mutational frequency in key genes from whole population sequencing data.**

This plot highlights the frequency of mutations (including nSNPs and indels) in specific genes that appeared in week 8 and persisted through weeks 15 or 24. A red cross indicates insufficient sequencing reads for the corresponding species and week, leading to no data analysis for that point.

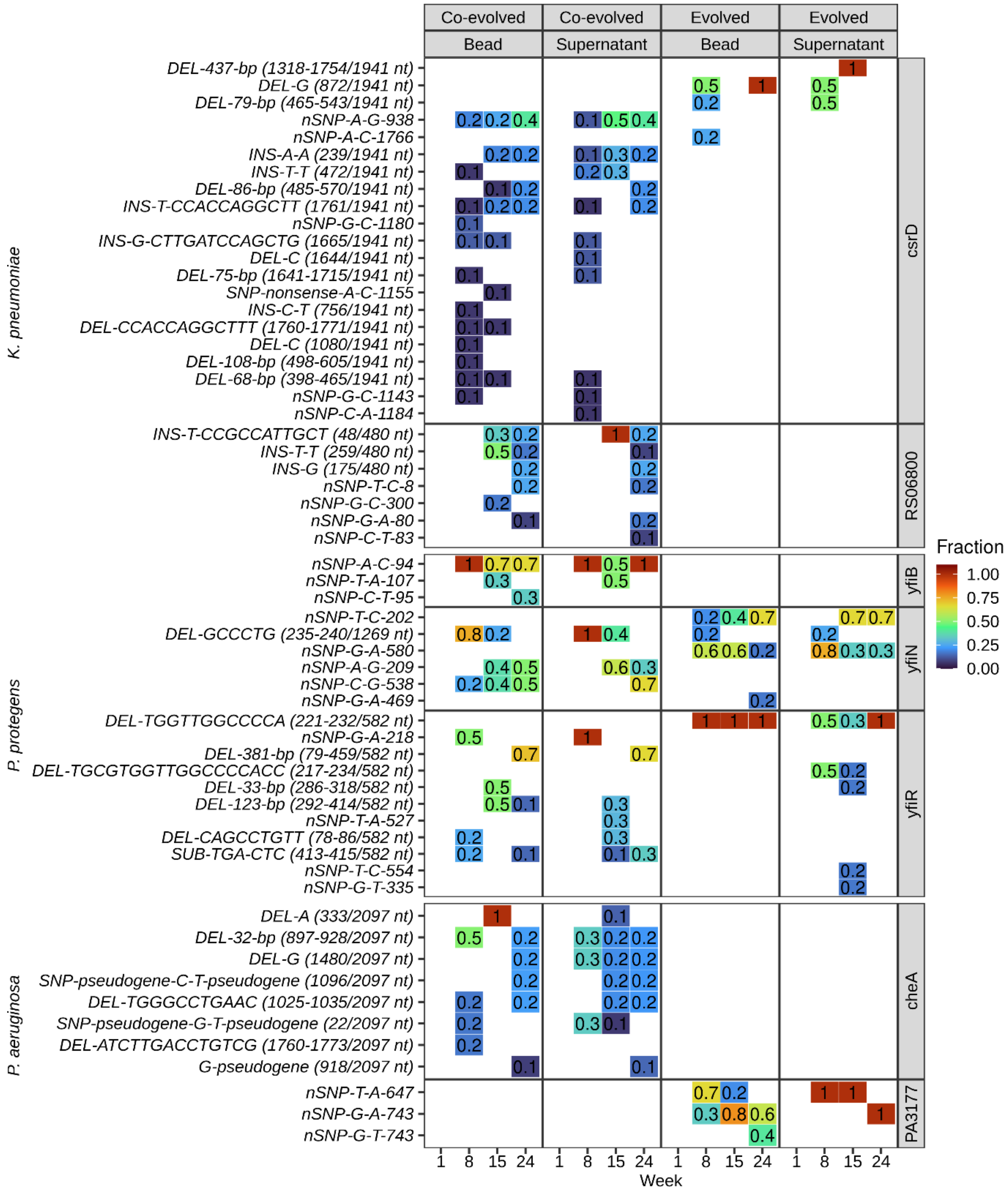


**Supplementary Fig. S4: Mutational spectrum in key genes of *K. pneumoniae*, *P. protegens* and *P. aeruginosa*.**

This heatmap illustrates the spectrum of mutations (nSNPs and indels) in key genes. The color and numerical values within the heatmap represent the fraction of isolates carrying a mutation in the corresponding gene.
